# Brain and circulating steroids in an electric fish: relevance for non-breeding aggression

**DOI:** 10.1101/2023.07.20.549867

**Authors:** Lucia Zubizarreta, Cecilia Jalabert, Ana C. Silva, Kiran K. Soma, Laura Quintana

**Author notes:** Current Address (Cecilia Jalabert). Indicates co-first author (Lucia Zubizarreta, Cecilia Jalabert). Indicates co-corresponding author.

## Abstract

Steroids play a crucial role in modulating brain and behavior. While traditionally it is considered that the brain is a target of peripheral hormones produced in endocrine glands, it has been discovered that the brain itself produces steroids, known as neurosteroids. Neurosteroids can be produced in brain regions involved in the regulation of social behaviors and can act locally regulating behaviors like reproduction and aggression. Here, for the first time in a teleost fish, we used liquid chromatography-tandem mass spectrometry (LC-MS/MS) to quantify a panel of 8 steroids in both plasma and brain. We use the weakly electric fish *Gymnotus omarorum*, a species which shows non-breeding aggression in both sexes, to characterize these hormonal profiles in wild non-breeding adults. We show that: 1) systemic steroids in the non-breeding season are similar in both sexes, although only males have circulating 11-KT, 2) brain steroid levels are sexually dimorphic, as females display higher levels of AE, T and E1, and only males had 11-KT, 3) systemic androgens such as AE and T in the non-breeding season are potential precursors for neuroestrogen synthesis, and 4) estrogens, which play a key role in non-breeding aggression, are detectable in the brain (but not the plasma) in both sexes. These data fall in line with previous reports in *G. omarorum* which show that non-breeding aggression is dependent on the estrogenic pathway, as has also been shown in bird and mammal models. Overall, our results constitute a fundamental groundwork to understanding the complexity of hormonal modulation, its potential sex differences, the role of neurosteroids and the interplay between central and peripheral hormones in the regulation of behaviors.

## Introduction

Steroids are potent signaling molecules that modulate brain and behavior. Historically, the brain has been considered a target for steroids produced in peripheral endocrine glands. However, the identification of steroidogenic enzymes in the brain reveals that the brain is also a source of steroids (reviewed in ^1, 2^). Neurally synthesized steroids, ‘neurosteroids’, play a wide variety of roles, such as the maintenance of synaptic transmission and connectivity in the hippocampus (reviewed in ^3^), and the regulation of neuroinflammatory responses (reviewed in ^4^), neuroprotective processes ^5^) and pain^6^. In addition, in both birds and mammals, neurosteroids, neuroestrogens in particular, are important in the regulation of social behaviors, including reproduction^7–9^ and aggression^10, 11^.

Several studies have quantified steroid levels in specific brain regions involved in natural behaviors. As an example, in zebra finches, brain microdialysis and immunoassay techniques have shown that in the auditory cortex of females a male song causes a transient increase of local androgens, with no effect on circulating androgens. These neuroandrogens, which can be aromatized into neuroestrogens, influence song processing and sensorimotor integration^12^. The use of ultrasensitive quantification techniques that can measure multiple steroids at low analyte levels can further our understanding of local steroidogenesis, the precursors involved, and the balance between local and systemic signaling^13, 14^. In wild caught song sparrows, *Melospiza melodia*, a comprehensive picture of the steroidal environment in the circulation and brain was achieved for the first time by performing an extensive profiling of 10 steroids in 10 areas of the social behavioral network by liquid chromatography-tandem mass spectrometry (LC-MS/MS). This study showed seasonal patterns in different regions of the social brain that differ greatly from the circulating values^13^.

Sex steroids are key regulators of aggression, an adaptive behavior that is displayed across species in different contexts. Although aggression is mostly studied in reproductive contexts, many species display this behavior outside of the breeding season, when circulating levels of sex steroids are often very low. These species offer an opportunity to study neuromodulation of aggression independently from gonadal steroid secretion^11, 15–18^. Song sparrows maintain high male-male territorial aggression even during the non-breeding season, when neuroestrogens are key in sustaining this behavior^11, 19–25^, similar to some rodents^10, 26^. These neuroendocrine mechanisms supporting aggression may be a common strategy across vertebrates^27, 28^.

Males and females of the weakly electric fish, *Gymnotus omarorum* ^29^ display high levels of aggression in the non-breeding season^30, 31^. In this species, castration does not affect aggressive behavior in non-breeding males. However, inhibition of aromatase reduces non-breeding aggression in both males and females, indicating that extra-gonadal estrogens are key to sustaining non-breeding aggression^18, 31^. Moreover, a transcriptomic analysis of the forebrain shows that dominant non-breeding male *G. omarorum* have a steroidogenic pathway directed towards estrogen synthesis whereas subordinate males have a pathway directed towards the production of non-aromatizable androgens^32^. These results support the hypothesis that brain-derived estrogens play an important role in the regulation of non-breeding aggression. However, no attempts have been made so far to measure blood and brain levels of sex steroids to reveal whether there is neural synthesis of key steroids, and identify sex differences.

Here, for the first time in a teleost fish, we used LC-MS/MS to quantify a panel of 8 steroids, including progesterone, cortisol, dehydroepiandrosterone (DHEA), androstenedione (AE), testosterone (T), 11-ketotestosterone (11-KT), estrone (E_1_), and 17β-estradiol (E_2_), in both plasma and brain in wild non-breeding male and female *G. omarorum*. Then, we examined sex differences in plasma and forebrain steroid levels. Finally, we evaluated the role of the brain as a potential source of estrogens and other sex steroids.

## Materials and Methods

### Field procedures

Free-living adult male (n=11) and female (n=11) *G. omarorum* were captured in the non-breeding season (June 2018) in Laguna de los Cisnes, Maldonado, Uruguay (34° 48’ S, 55° 180’ W). Sample collection was carried out during daytime, which corresponds to the resting phase of the animals. The capture method consists of locating the individuals with a detector without disturbing them and then using a rigid net to quickly lift the vegetation and the fish. Individuals were anesthetized immediately after capture by immersion in a fast-acting eugenol solution (1.2 mg/l). Blood was extracted from the caudal vein with a heparinized syringe, and body length was measured. Subjects were rapidly decapitated, and the brain was removed from the skull, quickly frozen in powdered dry ice, and stored in dry ice until arrival at the laboratory. To prevent a stress response from affecting basal steroid levels, there was a maximum of 3 min between capture and decapitation^33–35^. The time between decapitation and brain freezing was always less than 90 sec. Blood was kept on wet ice until centrifugation in the laboratory (approximately 5 h). Blood and brain samples that exceeded the stipulated times were excluded from the study, and were used for method development. The gonads were then removed and stored on ice. Once in the laboratory, gonad and body weights were determined. To calculate the gonadosomatic index, the weight of the gonads was added to the body weight, and the index was calculated as follows: [gonad weight / body weight] × 100. The blood was centrifuged (14,000g, 10 min) and plasma was collected and stored at -80, along with the brain.

All research procedures complied with ASAP/ABS Guidelines for the Use of Animals in Research and were approved by the Institutional Ethical Committee (Comisión de Ética en el Uso de Animales, Instituto Clemente Estable, MEC, CEUA-IIBCE: 001/02/2018).

### Brain dissection

The dissection of forebrains was carried out on a frozen metal plate surrounded by dry ice, under a magnifying glass. In each individual we obtained a brain section of the forebrain and part of the midbrain excluding the dorsally located cerebellum and optic tectum, as these regions do not have a major role in the regulation of social behaviors. To do so, frozen brains were mounted in tissue-tek, and fixed to a petri dish by their dorsal surface, exposing the ventral surface. We performed a coronal section with a heated cryostat blade immediately caudal to the inferior lobe (following ^36^). Then the section was remounted on tissue-tek to remove the cerebellum, optic tectum and in some cases in which the pituitary gland was still attached, we removed it as well. The dissected forebrains were weighed, placed in 2 ml polypropylene vials (Sarstedt AG and Co.) and stored at -80°C until processing.

### Steroid extraction

Extraction was performed using liquid-liquid extraction followed by solid phase extraction (based on^37^). Steroids were extracted from 25μL plasma and ∼50mg brain tissue. Samples were placed in 2-mL polypropylene vials (Sarstedt AG & Co., Nümbrecht,Germany) containing five zirconium ceramic oxide beads (1.4-mm diameter, Fisher Scientific). Then 50 μL of the deuterated internal standards (progesterone-d9, cortisol-d4, DHEA-d6, testosterone-d5, E_2_-d4; C/D/N Isotopes Inc., Pointe-Claire, Canada) in 50% HPLC-grade methanol were added to each sample, standards, and water blanks (except double blanks) to track recovery and matrix interference for each sample. Then, 1 mL of GC-grade ethyl acetate (EA) was added to each vial, and samples were homogenized using a bead mill homogenizer at 4 m/s for 30 s (Omni International Inc., Kennesaw, GA). Samples were then centrifuged at 16,100g for 5 min, and 1 mL of supernatant was transferred to a borosilicate glass culture tube pre-cleaned with HPLC-grade methanol (VWR International). Then 1mL of EA was added to the remaining sample (to maximize extraction efficiency), homogenized, and centrifuged as before, and again 1 mL of supernatant was collected and combined with the initial EA. Then 0.5 mL of Mili-Q water was added, and samples were vortexed and centrifuged at 3200g for 2 min. The water was removed and discarded, and the EA was dried in a vacuum centrifuge at 60°C for 45 min (ThermoElectron SPD111V). The pellets were reconstituted with HPLC-grade methanol and then extracted using solid phase extraction ^37^. Plasma, standards, and blanks were reconstituted in 0.5 mL of methanol, whereas brain samples were reconstituted in 1 mL of methanol and only 0.5 mL were further used to avoid matrix effects (internal standards correct for sample reduction). Columns (C18, Agilent, Santa Clara, CA; catalog no. 12113045) were previously conditioned with 3 mL HPLC-grade hexane and then 3 mL HPLC-grade acetone and equilibrated with 3 mL HPLC-grade methanol. Steroid extracts were then loaded onto the column (0.5 mL per sample), eluted with 2 mL HPLC-grade methanol, and eluates were collected. Samples were vacuum dried as previously stated. Dried pellets were reconstituted with 55 μL of 25% HPLC-grade methanol in MilliQ water, transferred to 0.6 mL polypropylene microcentrifuge tubes (Fisher Scientific), and centrifuged at 16,100g for 2min. Then 50 μL of supernatant were transferred to a LC vial insert (Agilent, Santa Clara, CA) and stored overnight at -20C until injection.

### Steroid analysis by LC-MS/MS

Steroids were quantified using a Sciex QTRAP 6500 UHPLC-MS/MS system as previously described^37^. Samples were transferred into a refrigerated autoinjector (15°C). Then, 45 μL of resuspended sample were injected into a Nexera X2 UHPLC system (Shimadzu Corp., Kyoto, Japan), passed through a KrudKatcher ULTRA HPLC In-Line Filter (Phenomenex, Torrance, CA) followed by a Poroshell 120 HPH C18 guard column (2.1 mm) and separated on a Poroshell 120 HPH C18 column (2.1 × 50 mm; 2.7 μm; at 40°C) using 0.1 mM ammonium fluoride in MilliQ water as mobile phase A (MPA) and HPLC-grade methanol as mobile phase B (MPB). The flow rate was 0.4 mL/min. During loading, MPB was at 10% for 0.5 min, from 0.6 to 4 min the gradient profile was at 42% MPB, which was ramped to 60% MPB until 9.4 min. From 9.4 to 9.5 min the gradient was 60-70% MPB, which was ramped to 98% MPB until 11.9 min and finally a column wash from 11.9 to 13.4 min at 98% MPB. The MPB was then returned to starting conditions of 10% MPB for 1 min. Total run time was 14.9 min. The needle was rinsed externally before and after each sample injection with 100% isopropanol.

We used 2 multiple reaction monitoring (MRM) transitions for each steroid and 1 MRM transition for each deuterated internal standard (Table 1). Steroid concentrations were acquired on a Sciex 6500 Qtrap triple quadrupole tandem mass spectrometer (Sciex LLC, Framingham, MA) in positive electrospray ionization mode for all steroids except E_1_ and E_2_, which were acquired in negative electrospray ionization mode.

**Table 1.**
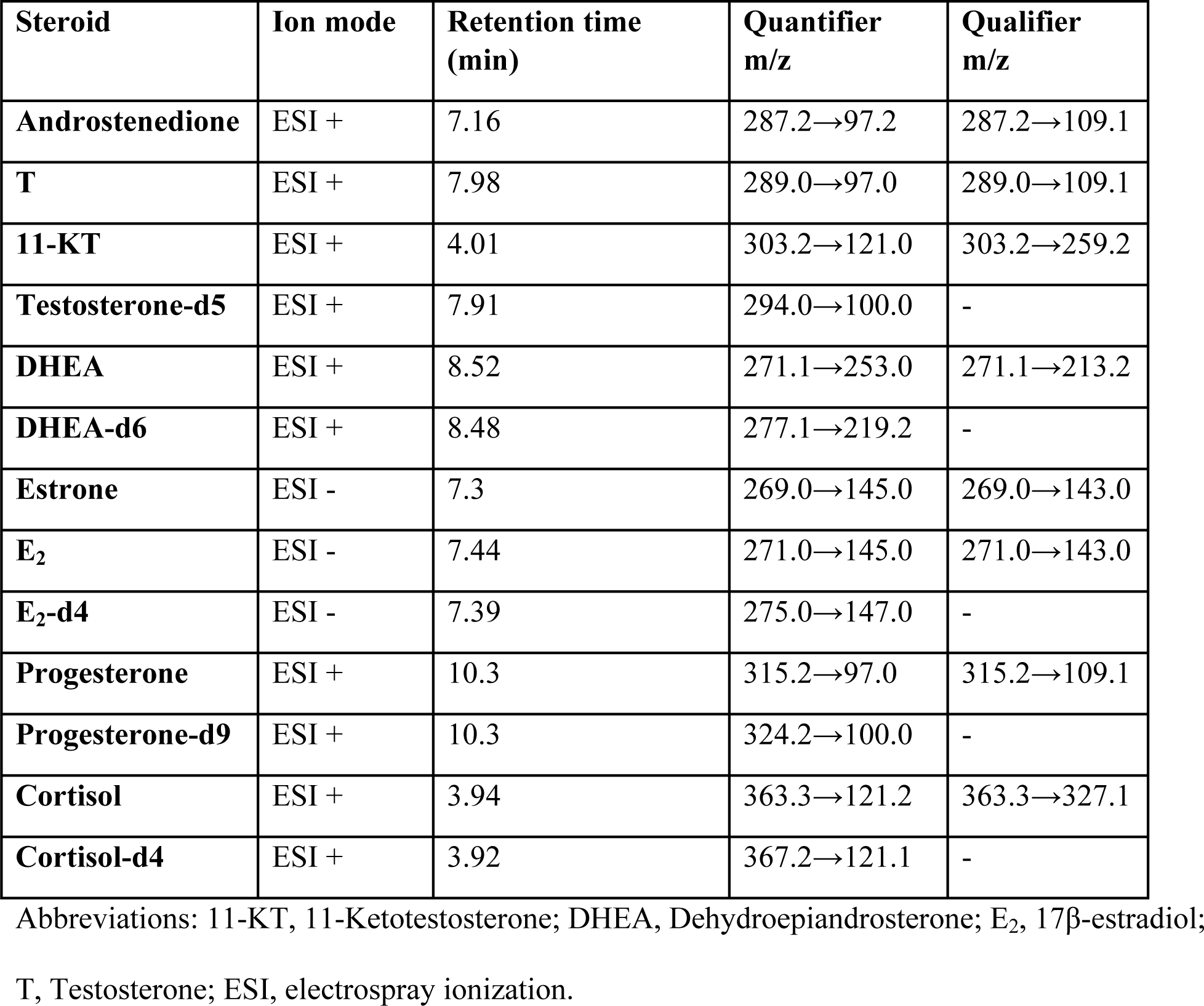
Scheduled multiple reaction monitoring for LC-MS/MS.

Calibration curves were made from certified reference standards (Cerilliant Co., Round Rock, TX) prepared in 50% HPLC-grade methanol. The calibration curve range was 0.2 to 1000 pg/tube for T and E_1_; 0.4 to 1000 pg/tube for AE and 11-KT; 0.8 to 1000 pg/tube for progesterone and E_2_; 2 to 1000 pg/tube for cortisol, and 20 to 10,000 pg/tube for DHEA (Table 2). Lower limit of quantification was calculated as the lowest standard on the calibration curve divided by the amount of sample (25µL for plasma and 25mg for brain) (Table 2). All blanks were below the lowest standard on the calibration curves.

**Table 2.**
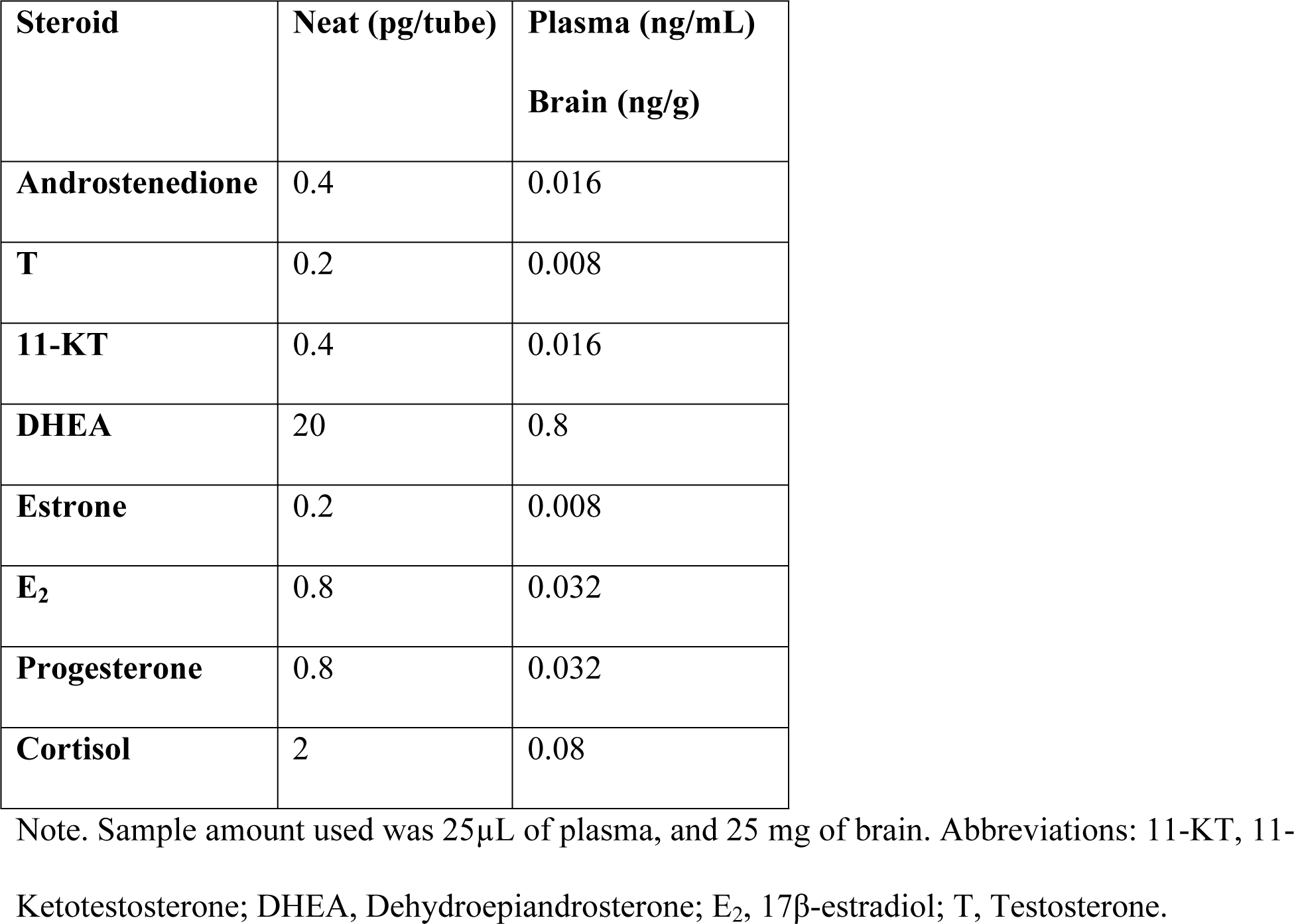
Lower limit of quantification in different matrices.

Matrix effects were assessed by extracting various amounts of sample (5, 15, 25, 50µL for plasma; and 2, 10, 20 mg for brain, n=3 per sample amount). For each sample, we measured the internal standard peak area and compared it with those in neat solution. Recovery was assessed using plasma and brain pools and comparing unspiked samples with samples spiked with a known amount of steroid (n=5 replicates per sample type). Accuracy was assessed by measuring quality controls with a known concentration in neat solution. Precision was evaluated by comparing replicates of quality controls within runs (intra-assay variation) and across runs (inter-assay variation). A total of 5 quality control replicates were run in each assay.

### Statistical analysis

A value was considered non-detectable if it was below the lowest standard on the calibration curve. When detectable samples in a group were ≥ 20%, missing values were estimated via quantile regression imputation of left-censored missing data using MetImp web tool^13, 38–42^. When detectable samples in a group were < 20%, non-detectable values were set to 0 to perform statistical analysis.

To compare steroid levels in the brain and circulation, and because the use of plasma overestimates steroid concentrations in the circulation^43–45^, we estimated steroid concentration in the blood. For this, we measured the hematocrit in adult male and female non-breeding *G. omarorum* (n=3). Plasma volume corresponds to 56% of blood volume and therefore steroid levels were multiplied by 0.56 to estimate steroid levels in whole blood.

Statistics were conducted using GraphPad Prism version 9.02 (GraphPad Software, La Jolla, CA, USA). When necessary, data were log transformed prior to analysis. When all values in a group were non-detectable in a brain region, 1 was added to each value of both groups and then data were log transformed. Sex differences in plasma and brain were analyzed by t-tests. To compare circulating and brain levels, data were analyzed using paired t-test (paired variables in the same fish). Correlations between steroid levels were examined using Spearman’s rho correlations with Benjamini-Hochberg correction for multiple comparisons (corrected p values are reported). Significance criterion was set at p ≤ 0.05. Graphs show the mean ± standard error of the mean (SEM) and are presented using the non-transformed data.

## Results

### LC-MS/MS assay development and validation

We developed a specific and sensitive method to quantify a panel of eight steroids by LC-MS/MS in both plasma and brain of *G. omarorum*. Analytes had distinct retention times and MRM transitions, which provide specificity (Table 1). Matrix effects, calculated by comparing internal standard peak areas of samples with those in neat solution (n=3/sample type), were similar for all steroids and within an acceptable range for plasma and brain tissue (Table 3). Recovery was assessed by subtracting unspiked sample pools from spiked sample pools and dividing by the amount of steroid added (n=5/sample type, Table 4). Recovery was within an acceptable range for most steroids in both plasma and brain tissue. Recovery for progesterone was over the acceptable limit (Table 4), suggesting that our assay overestimates progesterone levels; nonetheless, progesterone was non-detectable in plasma and brain tissue (see below). The assay demonstrated high accuracy and precision, with quality control measurements within the acceptable limits for accuracy and precision (Table 5).

**Table 3.** Matrix effects in biological samples.

| Steroid | Plasma<br>(% of peak area to neat) |  |  |  | Brain<br>(% of peak area to neat) |  |  |
| --- | --- | --- | --- | --- | --- | --- | --- |
|  | (5µL) | (15µL) | (25µL) | (50µL) | (2 mg) | (10 mg) | (20 mg) |
| **Testosterone-d5** | 137 | 124 | 84 | 77 | 93 | 98 | 100 |
| **DHEA-d6** | 66 | 88 | 96 | 84 | 90 | 102 | 87 |
| **E<sub>2</sub>-d4** | 79 | 71 | 67 | 73 | 113 | 98 | 96 |
| **Progesterone-d9** | 120 | 105 | 110 | 85 | 99 | 90 | 89 |
| **Cortisol-d4** | 89 | 92 | 106 | 96 | 107 | 96 | 100 |

Note. An increasing amount of samples were spiked with a fixed amount of steroid internal standard and the internal standard peak area was compared with those in neat solution. Samples were spiked with 2pg of testosterone-d5 and progesterone-d9, 20pg of Cortisol-d4 and E<sub>2</sub>-d4, and 60pg DHEA-d6. n=3 for each sample amount. Abbreviations: DHEA, Dehydroepiandrosterone; E<sub>2</sub>, 17β-estradiol.

**Table 4.** Recovery in biological samples.

| **Steroid** | **Plasma**<br>**(25µL)**<br>**recovery %** | **Brain**<br>**(25 mg)**<br>**recovery %** |
| --- | --- | --- |
| **Androstenedione** | 114 | 88 |
| **T** | 101 | 97 |
| **11-KT** | 135 | 93 |
| **DHEA** | 90 | 118 |
| **Estrone** | 86 | 96 |
| **E<sub>2</sub>** | 77 | 90 |
| **Progesterone** | 257 | 167 |
| **Cortisol** | 110 | 108 |

Note. Samples were spiked with 2 pg of each steroid except for DHEA (20 pg DHEA added), and cortisol (50 pg cortisol added). n=5 for each sample type. Abbreviations: DHEA, Dehydroepiandrosterone; E<sub>2</sub>, 17β-estradiol; T, Testosterone; 11-KT, 11-Ketotestosterone .

**Table 5.**
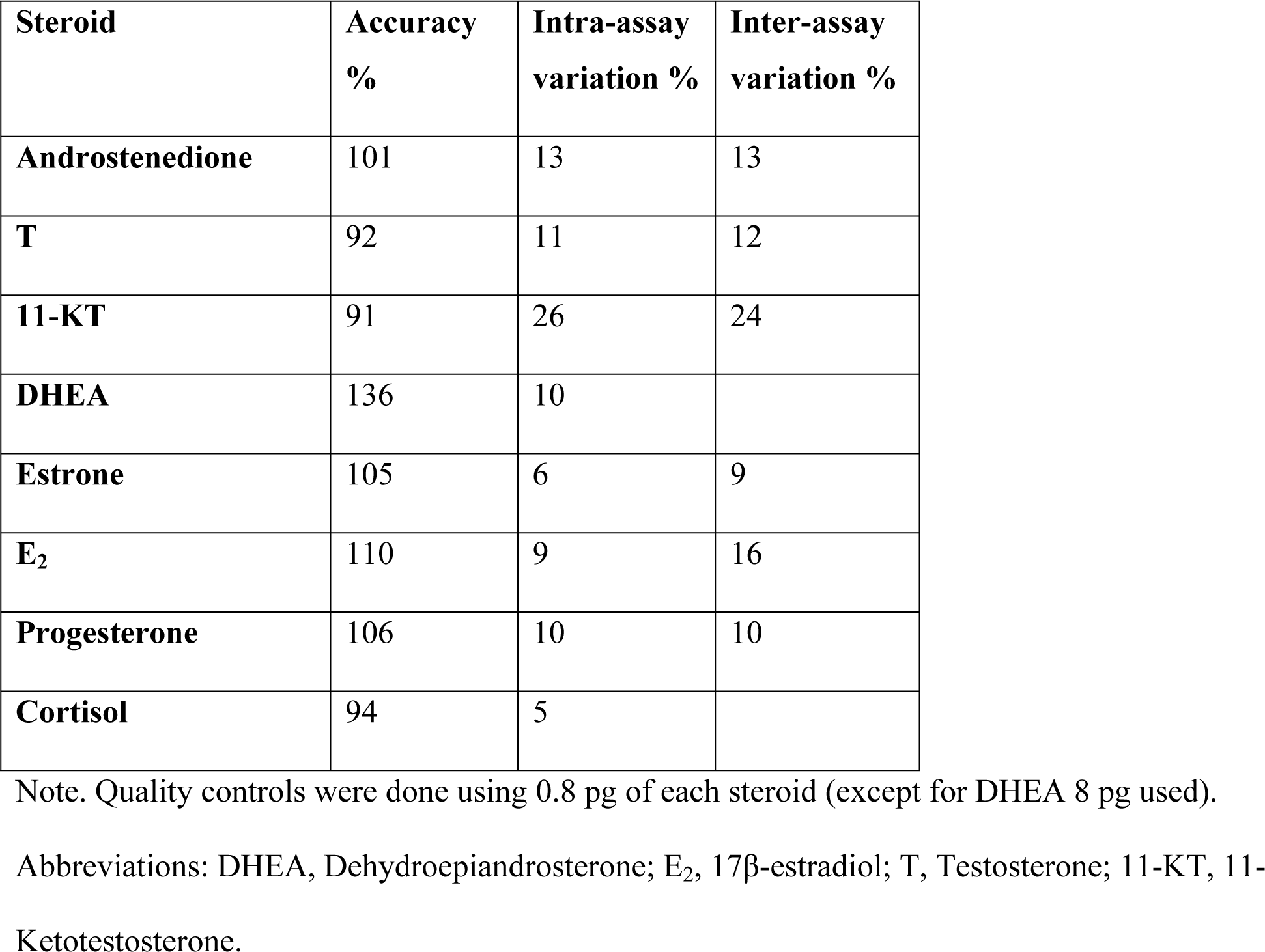
Assay accuracy and precision.

| **Steroid** | **Accuracy %** | **Intra-assay variation %** | **Inter-assay variation %** |
| --- | --- | --- | --- |
| **Androstenedione** | 101 | 13 | 13 |
| **T** | 92 | 11 | 12 |
| **11-KT** | 91 | 26 | 24 |
| **DHEA** | 136 | 10 |  |
| **Estrone** | 105 | 6 | 9 |
| **E<sub>2</sub>** | 110 | 9 | 16 |
| **Progesterone** | 106 | 10 | 10 |
| **Cortisol** | 94 | 5 |  |

Note. Quality controls were done using 0.8 pg of each steroid (except for DHEA 8 pg used).
Abbreviations: DHEA, Dehydroepiandrosterone; E<sub>2</sub>, 17β-estradiol; T, Testosterone; 11-KT, 11- Ketotestosterone.

### Systemic steroid levels

As shown in Fig 1A, non-breeding males had 4 detectable steroids in plasma: AE (0.14 ± 0.03 ng/ml), T (0.12 ± 0.02 ng/ml), 11-KT (0.07± 0.02 ng/ml), and cortisol (3.70 ± 0.95 ng/ml). On the other hand, females did not have detectable plasma 11-KT, but showed detectable levels of AE (0.15 ± 0.04 ng/ml), T (0.15 ± 0.04 ng/ml), and cortisol (5.2 ± 2.1 ng/ml). Circulating levels of steroids were not different between sexes (AE: t = 0.11, p = 0.90; T: t = 0.08, p = 0.93; and cortisol: t = 0.45, p = 0.66; Fig 1A) except for 11-KT (t = 4.3, p = 0.0005).

**Fig 1.**
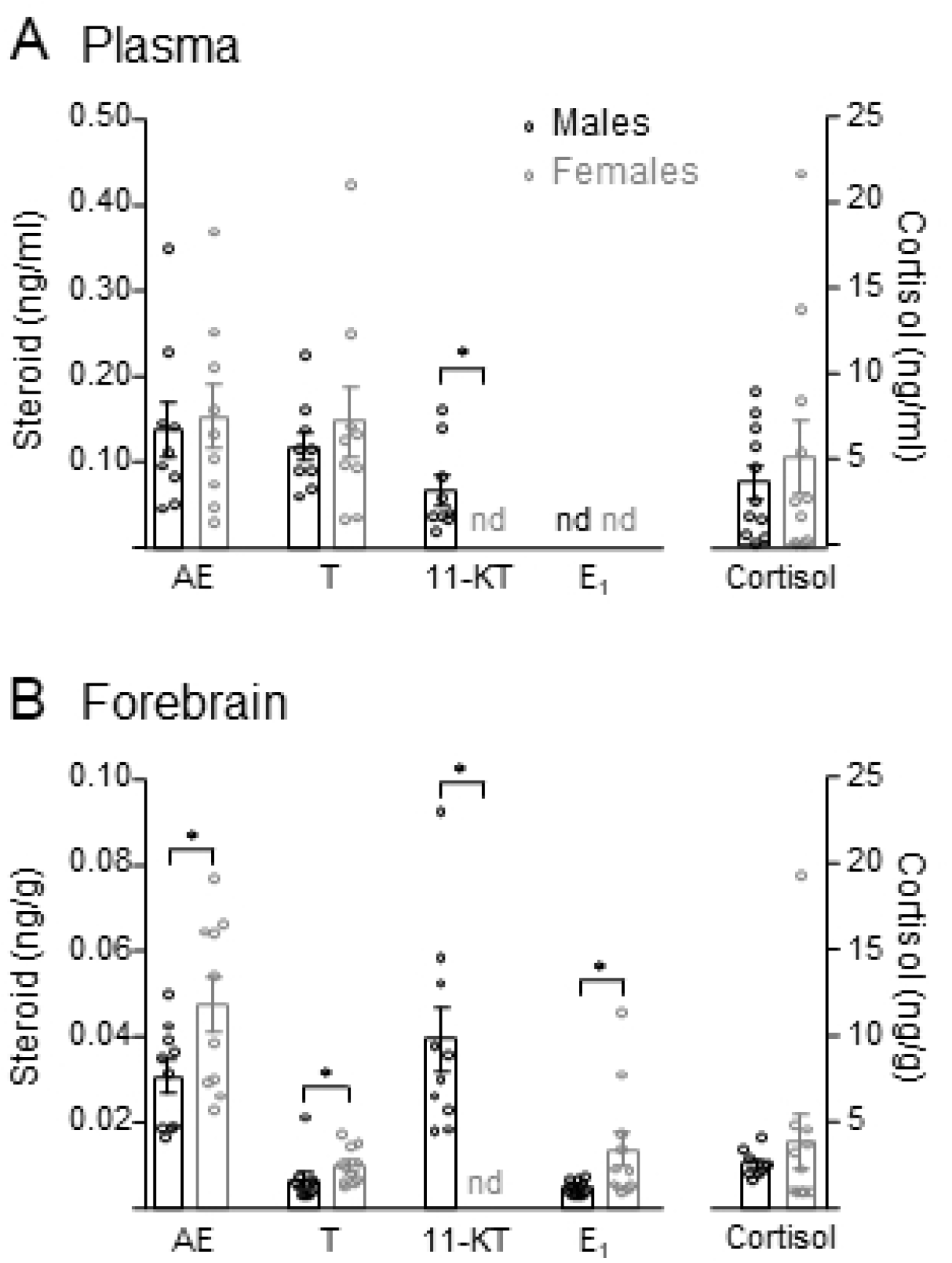
Steroid profiles of males and females of *G. omarorum* in the non-breeding season. Bar graphs show concentrations in plasma (A) and forebrain (B) expressed as the mean ± SEM, n = 11 per group. Abbreviations: AE, androstenedione; T, testosterone; 11-KT, 11-ketotestosterone; E1, estrone; nd, non-detectable. * p ≤ 0.05, *** p ≤ 0.001, **** p ≤ 0.0001.

Neither females nor males had detectable circulating levels of progesterone, DHEA, E_2_ and E_1_.

### Brain steroid levels

Forebrain steroid profiling in non-breeding males and females showed a similar pattern to plasma for androgens and cortisol, while estrogens were only detectable in the forebrain but not in plasma (Fig 1B). Males showed detectable forebrain levels of AE (0.03 ± 0.004 ng/g), T (0.06 ± 0.002 ng/g), 11-KT (0.04± 0.007 ng/g), and cortisol (2.54 ± 0.3 ng/g). Females showed AE (0.05 ± 0.006 ng/g), T (0.01 ± 0.001 ng/g), and cortisol (3.83 ± 1.6 ng/g) in the forebrain (Fig 1B). Both males and females showed detectable levels of forebrain E_1_ (males: 0.005 ± 0.0005 ng/g; females: 0.014 ± 0.004 ng/g), which was non-detectable in plasma samples (Fig 1A). Females had higher forebrain levels of AE (t = 2.152, p = 0.045), T (t = 2.246, p = 0.038), and E_1_ (t = 2.805, p = 0.01) than males. In contrast, 11-KT was higher in males than females (t = 5.3, p < 0.0001) as it was only detectable consistently in males (Fig 1B). Cortisol did not show a sex difference (t = 0.243, p = 0.81). Neither females nor males showed detectable forebrain levels of progesterone, DHEA, or E_2_.

### Comparison between circulating and brain levels

We compared brain and blood steroid levels for each subject by paired comparisons (Fig. 2). The analysis showed similar results in both sexes. Estrone was detectable in forebrain samples but not in plasma samples (males: t = 9.19; p < 0.0001; females: t = 3.35; p = 0.007; Fig 2). Androstenedione and T levels were higher in the blood than in the forebrain in both sexes (males: AE, t =3.05; p = 0.02; females: t = 2.15; p = 0.07; males: T, t = 13.35; p < 0.0001; females: T, t = 8.83; p < 0.0001; Fig 2). The analysis for 11-KT in males showed that blood and forebrain levels were not significantly different (t = 0.86; p = 0.42 Fig 2).

**Fig 2.**
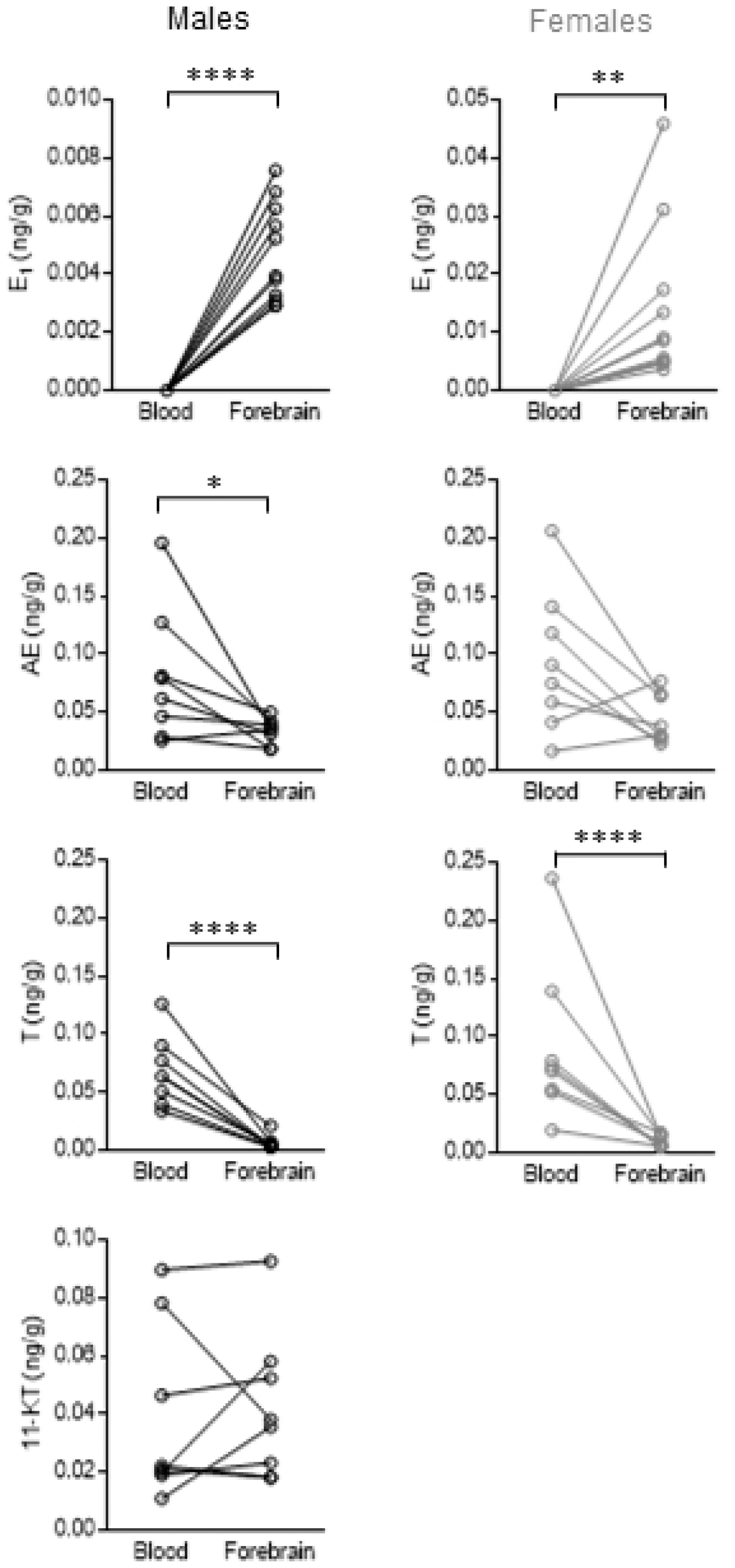
Comparison of circulating and brain steroid levels in *G. omarorum* in the non-breeding season. Blood and forebrain steroid concentrations were compared using a paired t-test. Lines connect values for the same fish. Abbreviations: AE, androstenedione; T, testosterone; 11-KT, 11-ketotestosterone; E1, estrone. * p ≤ 0.05, ** p ≤ 0.01, **** p ≤ 0.0001.

### Steroid correlates in the circulation and the brain

We assessed the relationship between circulating and brain steroid levels using correlation matrices (Spearman’s with Benjamini-Hochberg correction) (Fig 3). In males, the three circulating androgens showed positive associations (AE and T in plasma, r = 0.76, p = 0.013; AE and 11-KT in plasma, r = 67, p = 0.008; and T and 11-KT in plasma, r = 0.76, p = 0.012).

**Fig 3.**
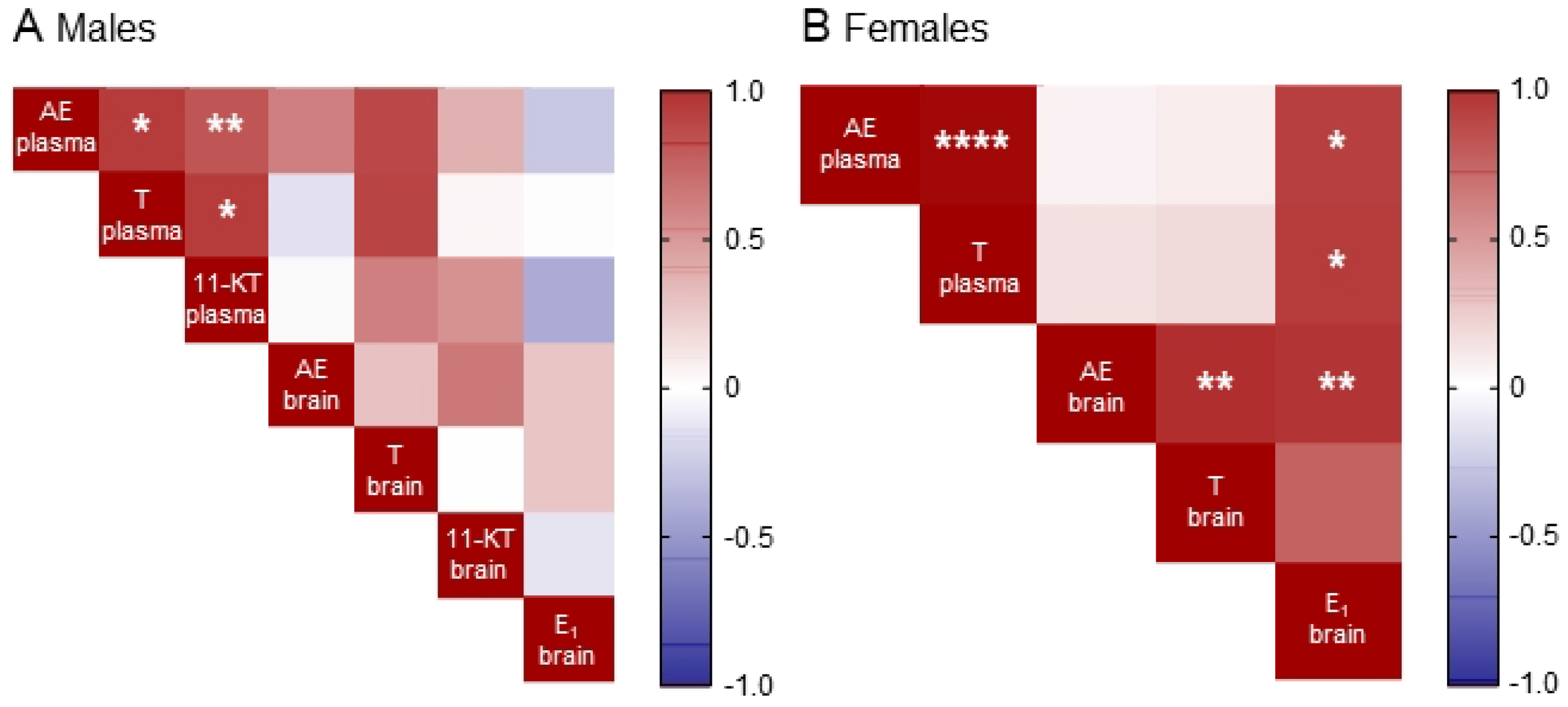
Correlation matrices between steroid levels in plasma and forebrain. Analyses were performed in males (A) and females (B). All values are expressed in Spearman’s rho and corrected for multiple comparisons using the Benjamini-Hochberg method. Abbreviations: AE, androstenedione; T, testosterone; 11-KT, 11-ketotestosterone; E1, estrone. * p ≤ 0.05, ** p ≤ 0.01, *** p ≤ 0.001, **** p ≤ 0.0001.

In females, circulating AE and T showed a positive association (r = 0.97, p < 0.0001). Brain T showed a positive association with brain AE (r = 0.82, p = 0.001). Brain E_1_ levels showed positive associations with two circulating androgens (brain E_1_ and plasma AE, r = 0.75, p = 0.05; brain E_1_ and plasma T, r = 0.77, p = 0.02), and a positive association with brain AE (r = 0.79, p = 0.006).

## Discussion

The present study is the first report of steroid profiling in the teleost fish brain using mass spectrometry. A panel of 8 steroids was validated for forebrain and plasma samples in male and female *G. omarorum*, a seasonal breeder in which both sexes display aggression in the non-breeding season. Here we show that: 1) systemic steroids in the non-breeding season are similar in both sexes, although only males have circulating 11-KT, 2) brain steroid levels are sexually dimorphic, as females display higher levels of AE, T and E_1_, and only males had 11-KT, 3) systemic androgens such as AE and T in the non-breeding season are potential precursors for neuroestrogen synthesis, 4) estrogens, which play a key role in non-breeding aggression, are detectable in the brain (but not the plasma) in both sexes. Taken together, these data provide fundamental insights into steroid regulation of behavior during the non-breeding season.

### Steroid measurement by mass spectrometry

The protocol developed in this study allowed us to describe brain and systemic steroid profiles for a teleost for the first time. The method combines liquid-liquid extraction with solid phase extraction to remove interference caused by the matrices, particularly challenging in lipid-rich brain samples. The LC-MS/MS assay has high specificity and high sensitivity (detection limits were generally 0.2 to 0.8 pg per sample). This is especially advantageous when quantifying steroids in non-breeding samples, as it allows for accurate quantification of low levels of analytes. In contrast, immunoassays often overestimate analyte levels (due to antibody cross-reactivity), especially at low analyte concentrations^46, 47^. Mass spectrometry also enables the measurement of steroids that are not commonly measured by immunoassays (e.g. AE and E_1_) and simultaneous quantification of multiple steroids in a sample. This allows a more comprehensive endocrine profile and a deeper understanding of systemic and local steroid levels in individual subjects.

### Plasma and brain steroid profiles and sex differences

There are few reports of brain steroid levels in non-breeding vertebrates in wild animals.

Resting, undisturbed non-breeding males and females had basal detectable circulating and forebrain levels of AE, T, and cortisol, while only males had circulating and forebrain detectable 11-KT. Systemic and forebrain levels of E_2_, DHEA, and progesterone were not detectable in either sex. In addition, E_1_ was detectable in both sexes exclusively in forebrain samples. The thorough systemic and brain steroid characterization achieved in this study is extremely valuable in the framework of understanding neural steroid synthesis and the hormonal mechanisms underlying social behaviors.

Circulating and forebrain cortisol levels in *G. omarorum* were at least an order of magnitude higher than the other steroids (Fig. 1A and B). Systemic glucocorticoid levels were comparable to those of non-breeding birds^13^ as well to those in non-breeding individuals of different teleost fish ^48^, although much lower than those of non-breeding mammals^49^. In non-breeding *G. omarorum* these levels did not present sex differences (Fig. 1A), as reported in other teleosts (reviewed in ^48^). Cortisol levels were high and similar between blood and brain (Fig. 1) suggesting that brain cortisol comes from a peripheral source.

Androgens were detectable both in circulation as well as in brain tissue. Circulating levels of AE and T were similar in males and females (Fig. 1A). However, the non-aromatizable androgen 11-KT, the main bioactive androgen of male teleosts^50^, showed the expected sex difference in plasma (Fig. 1A). Circulating 11-KT levels of non-breeding males were similar to those measured in this species by ELISA^18^ and to levels of other teleosts in the non-breeding season^51, 52^. In *G. omarorum*, forebrain AE and T were detectable in both sexes and higher in females than in males (Fig. 1B), in contrast to plasma, where no sex differences were observed (Fig. 1A). Males had forebrain 11-KT, which was undetectable in females. The only previous report of quantification of brain androgens in fish is in the bidirectional hermaphrodite *Lythrypnus dalli*, in which the authors explored the changes in 11-KT and T that accompany sex reversal, parental care and aggression^53–55^. In this species, brain T and 11-KT were present in both males and females^55^, most probably related to their particular sexual physiology. In non-breeding male song sparrows, basal levels of AE and T measured by LC-MS/MS were non-detectable^13^.

Overall, systemic steroid levels were similar between males and females, except for 11-KT, a typical male androgen (Fig. 1A). This lack of dimorphism was expected, as *G. omarorum* is a monomorphic species that shows no sex differences in body size, body condition, basal electric organ discharge^29^, spatial distribution in the field, or territory size^56^. Moreover, non-breeding territorial behavior shows no sex differences in dynamics, contest duration, outcome, or communication signals^57, 58^. Surprisingly, forebrain steroid levels did show sex differences (Fig. 1B), as females have higher levels of AE, T and E_1_ than males. This may be explained by: i) a sex difference in the expression of steroid binding globulins, which may lead to differences in brain uptake and/or retention of steroids^59, 60^, or ii) sex differences in brain steroidogenic enzymes that synthesize or metabolize steroids^61, 62^. Although brain sex differences are usually related to sex differences in behavior, they can also compensate for basal sex differences in physiological conditions and ultimately produce a monomorphic behavioral output^63^. In the breeding season there is evidence of sexually differentiated behavior in *G. omarorum* which may be associated with structural or functional brain differences. In *G. omarorum* males and females have different territory size determinants suggesting differential energetic requirements and associated foraging behavior^56^. Furthermore, males of the genus Gymnotus are reported to show paternal care^64, 65^. Seasonal changes in the brain may compensate for these differences, ultimately leading to monomorphic behaviors in the non-breeding season.

### Neuroestrogens and the mechanisms underlying non-breeding aggression

The presence of steroidogenic enzymes to produce steroids, even *de novo*, has been demonstrated in the brain of many teleost species (reviewed in^66^). In teleosts, it is well known that brain tissue can produce T, E_2_, and 11-KT^66–69^. However, in many of these experiments hormone precursors and co-factors are used in saturating concentrations. This may give us limited information on the brain as a steroid source in natural processes, as the activation of enzymes, concentration of precursors, and local metabolism are dynamic factors that influence local steroid levels.

An important role for circulating steroid hormones during the non-breeding season is to serve as precursors that reach the brain and are locally converted into hormones that are key in particular processes^70–72^. In that sense, the quantification of circulating DHEA was of particular interest as is an inactive androgen precursor that has been linked with the maintenance of non-breeding aggression in birds and mammals^27, 70^. However, circulating DHEA was non-detectable in *G. omarorum* during the non-breeding season indicating that it is absent or very low in the circulation similar to a recent report in song sparrows^13^. It has been reported in the circulation and in high levels in brain nuclei in non-breeding territorial song sparrows^13^, where it is involved in the regulation of aggression^14, 73, 74^. In non-breeding song sparrows progesterone is a possible precursor for neural synthesis of androgens and estrogens^13^. In *G. omarorum* progesterone was undetectable, although there are reports that teleost fish do have other progestogens^75^, which should be examined in future studies.

Neurally synthesized sex steroids are key in the regulation of social behavior. In zebra finch, quantification of brain steroids in the auditory cortex shows local synthesis of androgens in response to social stimuli, although androgens are also produced systemically^12^. In song sparrows, non-breeding aggression is not affected by androgen receptor antagonism^76^. However, in dusky gregories (a teleost fish), androgen receptor antagonism in the non-breeding season reduces aggression in males but not in females^51^. In the present study, androgens may be acting as precursors. Our results unequivocally show neuroestrogen synthesis in non-breeding females and males *G. omarorum*. In particular, E_1_ was detected in all forebrain samples in both sexes, while its plasma levels were non-detectable (Fig. 1 and Fig. 2). The correlations between hormonal precursors and their products provide insight into the pathways of steroid neurosynthesis. The direct precursor of E_1_ is AE, which is detectable in plasma and brain, although in much higher levels in circulation (Fig. 2). In females, there is a positive correlation between forebrain AE and E_1_, as well as between circulating AE and brain E_1_ (Fig. 3B). This suggests that peripheral AE is taken up by the brain and converted to E_1_, although we cannot rule out that there may be brain synthetized AE. Both AE and T are substrates of aromatase, and direct precursors to estrogens (E_1_ and E_2_ respectively). Nevertheless, brain AE concentrations are almost five-fold higher than those of T, and thus may be a stronger competitor for aromatase, accounting for the detection of E_1_ but not E_2_. The neurosynthesis of estrogens is consistent with the observation that in *G. omarorum* the dynamics and outcome of aggressive behavior depends on non-gonadal estrogen ^18, 31, 77^ and that there is differential expression of forebrain aromatase related to dominance and subordination in the context of non-breeding aggression^32^.

### Conclusions

Plasma and forebrain steroid profiles were characterized for the first time in non-breeding males and females of a teleost fish. Non-breeding individuals have detectable levels of circulating androgens and cortisol, brain androgens and cortisol, and brain E_1_. These results demonstrate the existence of estrogen neurosynthesis, both in females and males. The results of this study highlight the importance of characterizing circulating and brain steroid levels and are the starting point for the generation of hypotheses about the maintenance of non-breeding aggression and common neuroendocrine strategies across species.

## Acknowledgments

We thank Chunqi Ma for her fundamental assistance with protocol development in LC-MS/MS. Adriana Migliaro, Guillermo Valiño, and Rossana Perrone for help with fieldwork.

